# A Toolbox for Discrete Modelling of Cell Signalling Dynamics

**DOI:** 10.1101/249888

**Authors:** Yasmin Z. Paterson, David Shorthouse, Markus W. Pleijzier, Nir Piterman, Claus Bendtsen, Benjamin A. Hall, Jasmin Fisher

## Abstract

In an age where the volume of data regarding biological systems exceeds our ability to analyse it, many researchers are looking towards systems biology and computational modelling to help unravel the complexities of gene and protein regulatory networks. In particular, the use of discrete modelling allows generation of signalling networks in the absence of full quantitative descriptions of systems, which are necessary for ordinary differential equation (ODE) models. In order to make such techniques more accessible to mainstream researchers, tools such as the BioModelAnalyzer (BMA) have been developed to provide a user-friendly graphical interface for discrete modelling of biological systems. Here we use the BMA to build a library of discrete target functions of known canonical molecular interactions, translated from ordinary differential equations (ODEs). We then show that these BMA target functions can be used to reconstruct complex networks, which can correctly predict many known genetic perturbations. This new library supports the accessibility ethos behind the creation of BMA, providing a toolbox for the construction of complex cell signalling models without the need for extensive experience in computer programming or mathematical modelling, and allows for construction and simulation of complex biological systems with only small amounts of quantitative data.

**AUTHOR SUMMARY:** Ordinary differential equation (ODE) based models are a popular approach for modelling biological networks. A limitation of ODE models is that they require complete networks and detailed kinetic parameterisation. An alternative is the use of discrete, executable models, in which nodes are assigned discrete value ranges, and the relationship between them defined with simple mathematical operations. One tool for constructing such models is the BioModelAnalyzer (BMA), an open source and publicly available (https://www.biomodelanalyzer.org) software, aimed to be fully usable by researchers without extensive computational or mathematical experience. A fundamental question for executable models is whether the high level of abstraction substantially reduces expressivity relative to continuous approaches. Here, we present a canonical library of biological signalling motifs, initially defined by Tyson et al (2003), translated for the first time into the BMA. We show that; 1) these motifs are easily and fully translatable from continuous to discrete models, 2) Combining these motifs in a computationally naïve way generates a fully functional and predictive model of the yeast cell cycle.

## INTRODUCTION

We are in an era of ever-increasing biological data. With data available from genomic studies, through to metabolomic studies, the size, scale and heterogeneity of the resources available present many triumphs in terms of advancing high-throughput technologies but also many challenges. Despite the enormous multitude of available data, our understanding of how such information encoded in a cell’s genome is used to carry out the complex biological interactions found between genes and gene products is still lacking. It is therefore no surprise that a central goal of modern biology in this pos-tgenomic era is to understand the structural and temporal nature of these
control networks. Not only would this allow us to translate ‘Big Data’ into working models of biological systems, but also equip us with a better understanding of biological mechanisms, allowing the exploration of emergent behaviours and consequences of genomic variants, with an aim to develop real-world hypotheses for experimental validation.

If we are to meet these challenges, new tools, techniques and ways of working need to be adopted. Whilst experimental procedures using a traditional reductionist approach, focusing on the study of individual proteins or genes in isolation from other network interactions have proved useful in uncovering specific elemental functions of various cellular mechanisms, many disease processes continue to elude us. This has fuelled the growth of new lines of scientific inquiry. The wide-ranging, vast improvements in computing power brought about at the beginning of the twenty-first century has led biologists down the path of Systems Biology as a means to organise this biological data more holistically. This strategy therefore seeks to combine traditional biological thinking with more interdisciplinary, integrated, synthetic approaches allowing for larger-scale simulations of complex systems, which could revolutionise biomedical discovery.

Computational modelling therefore presents a powerful and novel approach to combat these challenges. The application of standard mathematical modelling, such as through stochastic or ordinary differential equations (ODEs), have been faithfully reproducing the interplay between genes and proteins in small regulatory networks with relative success. Prominent examples of ODE models include that of bacterial chemotaxis [1], the lactose operon control system in *Escherichia coli* [2] and the process of X chromosome inactivation [3], the cell cycle in yeast [4], and the generation of amyloid fibrils [5].Such models employ complex kinetic equations to describe relationships between proteins or genes over time, and require highly accurate and intensive experimental data for their development as input. The complexity of such equations and experimental data required can provide a lot of dynamical detail however this complexity also begs the question of whether this approach will scale well when constructing much larger, more intricate networks in the future.

Executable modelling on the other hand, which describes biological systems as discrete systems, can provide a much simpler class of models [6]. Such models are immediately executable, allowing for much larger-scale simulations to be produced as well as providing the ability to undergo model checking and other formal verifications with ease [7–10]. One of the oldest and simplest forms of executable network models is based on Boolean states (logical models), where each node of the network represents a single gene or protein which is in one of two states: active/on (1) or inactive/off (0) [11,12]. Abstract models based on this paradigm have proved capable of forecasting dynamic processes. Clear examples include that of the control of segment polarity genes in Drosophila [13] or modelling of the neurotransmitter-signalling pathway between dopamine and glutamate receptors [14]. Yet the activities of cellular networks and signalling pathways are often subtler than this, which has resulted in various extensions being made to this model. One such refinement is *Qualitative Networks* (QNs), which uses discrete variables as opposed to Boolean states, and is able to model a much broader range of interactions by using algebraic target functions [7]. These target functions are composed of simple mathematical operations (e.g. addition, subtraction, division, multiplication) to allow for the generation of models with complex relationships between variables.

The BioModelAnalyzer (BMA) tool is a freely accessible online platform that creates QNs from user’s instructions. These instructions are formed using a graphical interface, where different genes or proteins are represented by simple symbols that can be connected by inhibitory or activatory edges negating the need for extensive experience in computer programming, logical formalisms or mathematical proofs [15]. As a result, the BMA is a highly accessible, unimposing interface that is suitable for experimental biologists, whilst still providing powerful stability checking, simulation and Linear Temporal Logic analysis abilities. Although on the surface BMA may appear to be highly abstract, elaborate biological functions can be robustly modelled such as that of *C. elegans* germline development [16], mammalian epidermis differentiation [7], gene and protein regulatory networks in chronic myeloid leukaemia (CML) [17] and acute myeloid leukaemia (AML) [18]. In the case of CML, a novel therapeutic strategy using an Imatinib and pan-Bcl2 family gene inhibitor combination has been identified, highlighting BMAs ability to work on either a hypothesis-creation or hypothesis-testing basis. Cell line specific differences in the PIM pathway were identified in the case of AML, leading to clinically relevant predictions about resistance and how to overcome it.

Although BMA provides the ability to encode complex dependencies between different genes or proteins via the use of algebraic target functions, this task can still seem quite onerous to many biologists. In 2003, Tyson, Chen and Novak [19] published a review outlining a concise mathematical vocabulary of common cellular interactions and pathways using ODEs. In their article, they identify a number of simple functional motifs, akin to electrical circuits which are found at the base of a variety of key biological processes and can be easily combined in order to model complex regulatory interactions. Here we outline a target function library which translates the ODEs outlined by Tyson et al. [19] into discrete equations encoded within nodes of a BMA model. In order to investigate whether these target functions are capable of modelling cellular behaviours of greater complexity, we then created a BMA model of eukaryotic cell cycle regulation similar to Tyson et al [19]. *In silico* over-expression and knockouts of combinations of genes and genetic interactions were then carried out to highlight the sensitivity of our model. A key benefit of using discrete, executable modelling is that complex systems can be simulated and analysed, and experimentally testable hypotheses can be generated in the absence of large amounts of quantitative data required for ODE models.

This library of ODE translations to discrete target functions also complements the accessibility ethos behind the creation of the BMA. By providing simple building blocks that can be “plugged” into a set of specific nodes, much time and effort will be saved allowing biologists to construct elaborated valid models of biological phenomena, which can guide and direct hypotheses and ultimately drug treatments.

## RESULTS

### A Target Function Library Accurately Reproduces Expected Biological Behaviour in Simple Networks

We constructed QN models representing the ten major archetypal regulatory and signalling pathways. Networks were generated within the BMA, and signal/response curves compared to previous publications [19–23] for accuracy. Networks are represented by a series of nodes interconnected via activatory (i.e. generally increasing target node value), and inhibitory (generally decreasing target node value) relationships. Nodes in the system can contain values with a granularity of 5 (a range of 0-4), but are generally easily extrapolated to different system ranges. Full details are included in **Supplementary Table 1**, and all models are available in supplementary data.

### 1. Linear Response

A system where the signal-response is linear (i.e. an increasing signal gives a proportionally increasing response) can be accurately modelled using the default target function. A node with no specified target function will have its value calculated by:

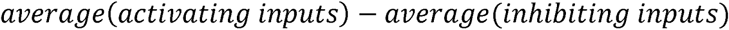

A schematic linear signal-response network, from Tyson et al. [19] and built within the BMA is shown in **Fig 1, A, i &ii**, with signal-response curves from both systems shown in **Fig 1, A, iii & iv.**

**Fig 1.**
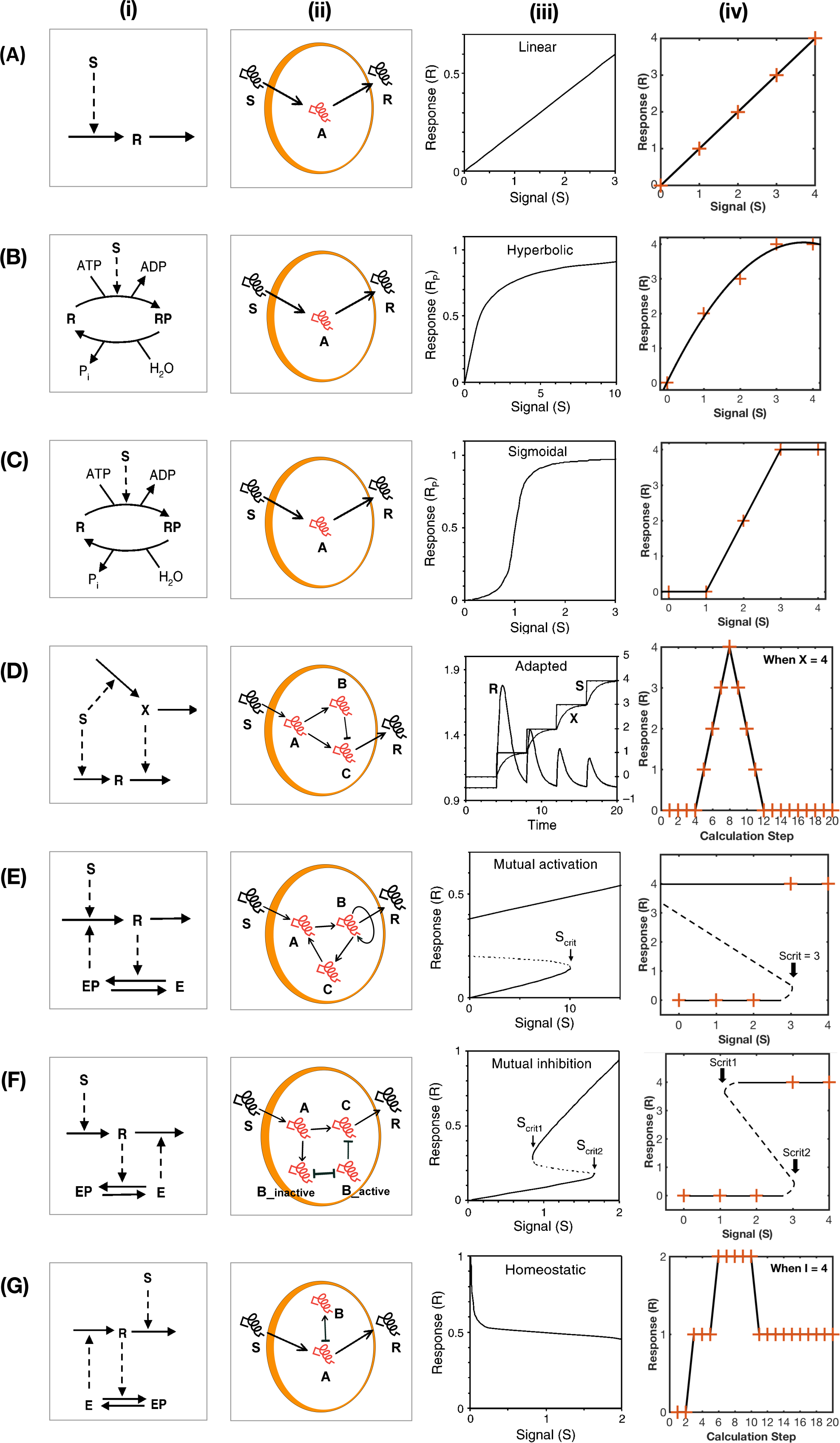
Comparison of Signal-Response Elements. In this illustration, the rows correspond to **(A)** linear response **(B)** hyperbolic response, **(C)** sigmoidal response, **(D)** perfect adaption, **(E)** mutual inhibition, **(F)** mutual inhibition and **(G)** homeostasis as in Tyson et al. [19] The columns correspond to **(i)** Tyson et al. [19] wiring diagrams, **(ii)** BMA wiring diagram translation, **(iii)** Tyson et al. [19] signal-response curves and **(iv)** BMA equivalent signal-response curves. Each BMA wiring diagram contains a unique set of target functions located within particular nodes of the network which can be found in **Supplementary table 1.** For most cases clear comparison between Tyson et al. [19] wiring diagrams **(i)** and the corresponding BMA wiring diagrams (ii) can be made. Here like in Tyson et al. [19] S indicates the input Signal and R indicates the output Response with, in our case, letters A-C representing intermediate nodes. The graphs in (iv) are derived from simulation analysis carried out in the BMA. For all cases bar (d- iv) and (g-iv) the signal is altered from 0 through to 4 directly within the S node and the output in node R recorded and subsequently plotted. For cases (d- iv) and (g-iv) a simulation is run with a set signal input of 4 as an example, and the response output from the BMA simulation plotted based on the response per calculation time step. Graphs plotted from the BMA model (iv) can then be compared to ODE counterpart (iii). In (e-iv) and (f-iv) the dashed line represents an unsteady state. In (e-iv) S_crit_, which is denoted x in our target function (**Supplementary table 1**) represents the signal input where a switch in steady states will occur. The motifs in each case reproduce the bifurcation as expected. Similarly, in (f-iv) S_crit1_ which is denoted y in our target function and S_crit2_ which is denoted ed z in our target function also correspond to the switch points in stable states.

### 2. Hyperbolic Response

We generated four ways to discretely model different hyperbolic functions within the BMA. This function describes a “phosphorylation and dephosphorylation” reaction and is modelled using a three-node wiring diagram shown in **Fig 1**, **B**, **ii**. A simple function included in node A results in a hyperbolic response as a result of a linearly increasing input. Node A contains the target function:

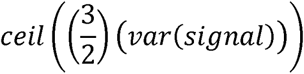

Where *var(signal)* represents the signal received by the network. This linear approximation captures the rapid initial growth of the response, whilst the plateau is enforced by the granularity of the variable. Additional modifiers (for details see **Supplementary Table 1**) can be included to change the shape and thresholds of the response. Hyperbolic signal-response curves from Tyson et al. [19] and from within the BMA are shown in **Fig 1**, **B**, **iii** **&** **iv**.

### 3. Sigmoidal Response

Sigmoidal response curves represent systems that act in a switch like manner, which are reversible and increase continuously with an increasing input. Schematics for sigmoidal signal-response networks are shown in **Fig 1**, **C**, **I** **&** **ii**. The target function for Node A contains the function describing the sigmoidal response. Multiple functions produce differing sigmoidal curves (for details see **Supplementary Table 1**). The simplest, producing a sigmoidal response from a linear signal is:

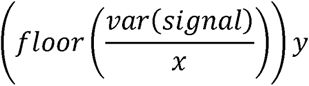

Where *x* is the value at which the system switches between high and low
values, and *y* is the upper value for the sigmoidal response. Signal-response curves are shown in **Fig 1**, **C**, **iii** **&** **iv**.

### 4. Perfect Adaptation Response

Adaptation is defined as “a process where a system initially responds to a stimulus, but then returns to basal or near-basal levels of activity after some period of time” [24]. Perfect adaptation is further characterised by the final response of the network returning to the exact pre-stimulus level. Perfect adaptation is used in numerous biological systems, for example, the Friedlander and Brenner [25] model of ion channel activation and inactivation. Network schematics for perfect adaptation systems can be seen in **Fig 1**, **D**, **i** **&** **ii**. Perfect adaptation is modelled with the addition of the following target function to Node C (Signal-response curves can be seen in **Fig 1**, **D**, **iii** **&** **iv**):

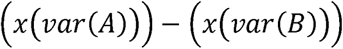

Where *x* represents the maximal height of the response before the system adapts.

### 5. Mutual Activation Response

Mutual activation behaviour represents irreversible cell switches, i.e. a “point-of-no-return”. These discontinuous, one-way switches are typical of cell fate determination. Once a critical signal value (S_crit_) is reached, the response immediately increases to a high level. A critical feature of mutual activation networks is that the switch is irreversible i.e. if the signal increases beyond S_crit_ and subsequently decreases, the response will not decrease. Network schematics for mutual activation systems are shown in **Fig 1**, **E**, **i** **&** **ii**. Inclusion of the following target function in Node A results in the signal-response curves shown in **Fig 1**, **E**, **iii** **&** **iv**:

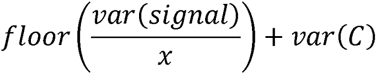

Where *x* is S_crit_ − the value at which the irreversible switch occurs.

### 6. Mutual Inhibition Signal Response Curve

Mutual inhibition differs from mutual activation in that these systems exhibit hysteresis; if the input decreases below a defined critical value, then the output will return to zero. Tyson et al. [19] describe this type of feedback as a “toggle switch”, where there are two defined critical values; S_crit1_ and S_crit2_, at which point the response will shift from either upper or lower values to the opposite. This is simplified below:

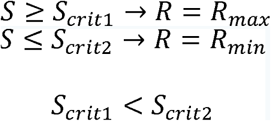

Essentially, this works similarly to mutual activation, except if S is decreased below Scrit2 then the switch will return to the inactive state. Our model is composed of 6 nodes and is compared to the “traditional” toggle switch schematic in **Fig 1, F, ii.** B is split into two separate nodes representing active and inactive states (B_active_, and B_inactive_ respectively), and it is the interactions between these 2 states of B that give rise to hysteresis. The target function for the node representing inactive B (Node B_inactive_) in a system with a granularity of 0-4 is:

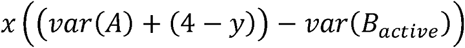

Where *x* represents the maximal response of the network, and *y* is S_crit1_. The target function for the node representing active B (Node B_active_) is:

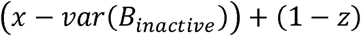

Where *x* represents the maximal response for the network, and *z* is S_crit2_. Additionally, Node C contains the following target function:

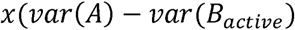

Where *x* represents the maximal response for the network. Signal-respons curves for this network are shown in **Fig 1**, **E**, **iii** **&** **iv**.

### 7. Homeostasis

Homeostatic regulation involves a network where the network counteracts the activity of the stimulus such that the response is constrained to a very narrow window (in the case of our network, a single value). A schematic homeostasis network is shown in **Fig 1**, **F**, **I** **&** **ii**. In this network, the granularity of Node B is adjusted such that it is double the range of the other nodes within the system. This is required for stability of the system, as nodes can only alter by a single integer value each time step, a node of granularity 10 (0-9) will take twice as long to reach a maximal value from 0 as a node with a granularity of 5 (0-4). This temporal difference allows us to eliminate instability in the system caused by oscillatory feedback between Nodes A and B, which is present when both nodes have the same granularity. Within this network two unique target functions are required to exhibit homeostasis, for Node A:

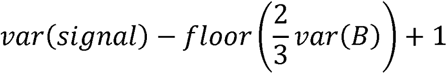

And for the node representing the system response (Node Response):

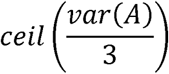

Signal-response curves can be seen in **Fig 1**, **F**, **iii** **&** **iv**.

### 8. Negative Feedback Oscillations

Negative feedback oscillations result from similar network wiring as homeostasis, with the result being a system where the response oscillates between 0 and the signal value.

A negative feedback oscillation loop is seen in **Fig 2**, **A**. No change in the default target functions are required to generate an oscillatory output. For these networks, an input of S will result in an oscillation that tends to between S and 0. The temporal constraints of the network however (in that each node can only update by a single integer value each step) results in cases where the oscillation will not reach the maximal value before the inhibitory portion of the network kicks in. The ultimate range of the oscillations can be tailored however, with either the addition of values to the output node (in order to adjust the oscillation range up or down), or by inserting the following formula into node A:

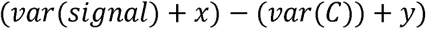

Where the difference between *x* and *y* changes the range of the oscillations. Additionally, the temporal properties of the system, specifically how long it takes to perform each loop, can be adjusted by the addition of more nodes to the loop, with a large number of nodes increasing the number of steps required to complete one oscillation. For an example see **Supplementary Figure 1.**

**Fig 2.**
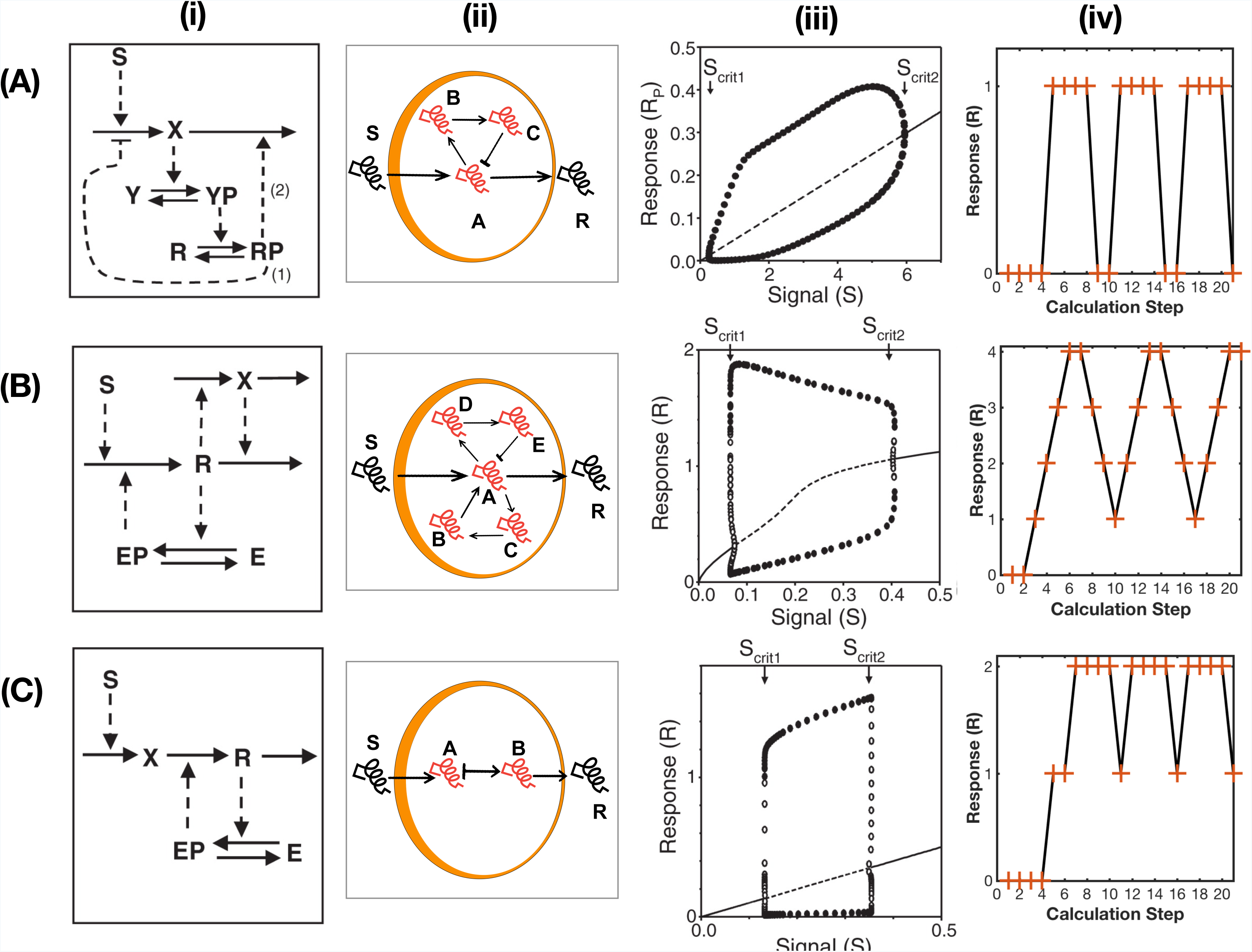
Comparison of Oscillatory Networks. In this illustration, the rows correspond to **(A)** negative feedback, **(B)** activator-inhibitor and **(C)** substrate-depletion oscillators as in Tyson et al. [19] The columns correspond to **(i)** Tyson et al. [19]wiring diagrams, **(ii)** BMA wiring diagram translation, **(iii)** Tyson et al. [19] signal-response curves and **(iv)** BMA equivalent signal-response curves. Each BMA wiring diagram contains a unique set of target functions located within particular nodes of the network which can be found in **Supplementary table 1.** For most cases clear comparison between Tyson et al. [19] wiring diagrams (i) and the corresponding BMA wiring diagrams (ii) can be made. Here like in Tyson et al. [19] S indicates the input Signal and R indicates the output Response with, in our case, letters A-E representing intermediate nodes. The graphs in (iv) are derived from simulation analysis carried out in the BMA. For all cases bar a simulation is run with a set signal input of 2 as an example, and the response output from the BMA simulation plotted based on the response per calculation time step and are thus not directly comparable, however clear oscillatory behaviour can still be observed.

### 9. Activator-Inhibitor Oscillations

The activator-inhibitor oscillation relationship is characterised by a positive and negative feedback loop within a system (shown in **Fig 2**, **B**, **i** and **ii**). The interactions of the two loops result in a system that oscillates between a maximal and minimal value, called a hysteresis oscillator. Including the following formula in node A results in an oscillation between the maximal and minimal values of the nodes when I = 2:

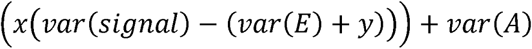

Where *x* is 3 value of the nodes, and *y* is 0. Adjusting these values will change the range of the oscillations (i.e a range of 3 or 2 is obtained by reducing the value of *x*), and altering the value of *y* adjusts the start and end points of the oscillation. The signal-response curves for activator-inhibitor networks are shown in **Fig 2**, **B**, **iii** and **iv**).

### 10. Substrate-Depletion Oscillations

The substrate depletion oscillation (SDOs) is quite similar to that of negative feedback. However, the number of nodes are reduced to reflect the greater intimacy between enzyme-substrate reactions compared to negative feedback loops. The network schematics for substrate-depletion oscillations are shown in **Fig 2**, **C**, **i**. In substrate-depletion oscillations, a small signal produces a small response and a large signal produces a large response. To model this, the following target function is applied to Node A:

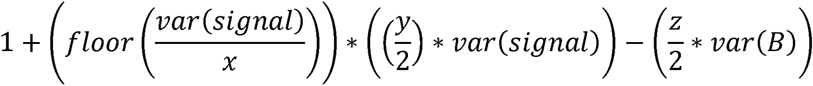

Where *x* is the starting point of the oscillations and *y* and *z* are the range of
the oscillations. Signal-response curves are presented in **Fig 2**, **C**, **ii**.

### Using the BMA Target Function Library to Construct Complex Networks: Eukaryotic Cell Cycle Control

After translating these motifs that were originally defined with ODEs to BMA models and their target functions, we then sought to determine the robustness of these motifs and their target functions by modelling a complex cellular behaviour: Eukaryotic cell cycle regulation. Based on the wiring diagram presented by Tyson et al. [19], a QN model was constructed using our own BMA target function library (**Fig 3**, **A**). Clear descriptions of the dynamics of cell cycle regulation can be found in the following review articles: Tyson, Csikasz-Nagy & Novak (2002) [22], Tyson & Novak (2015)[20] and Hochegger, Takeda & Hunt (2008) [26].

**Fig 3.**
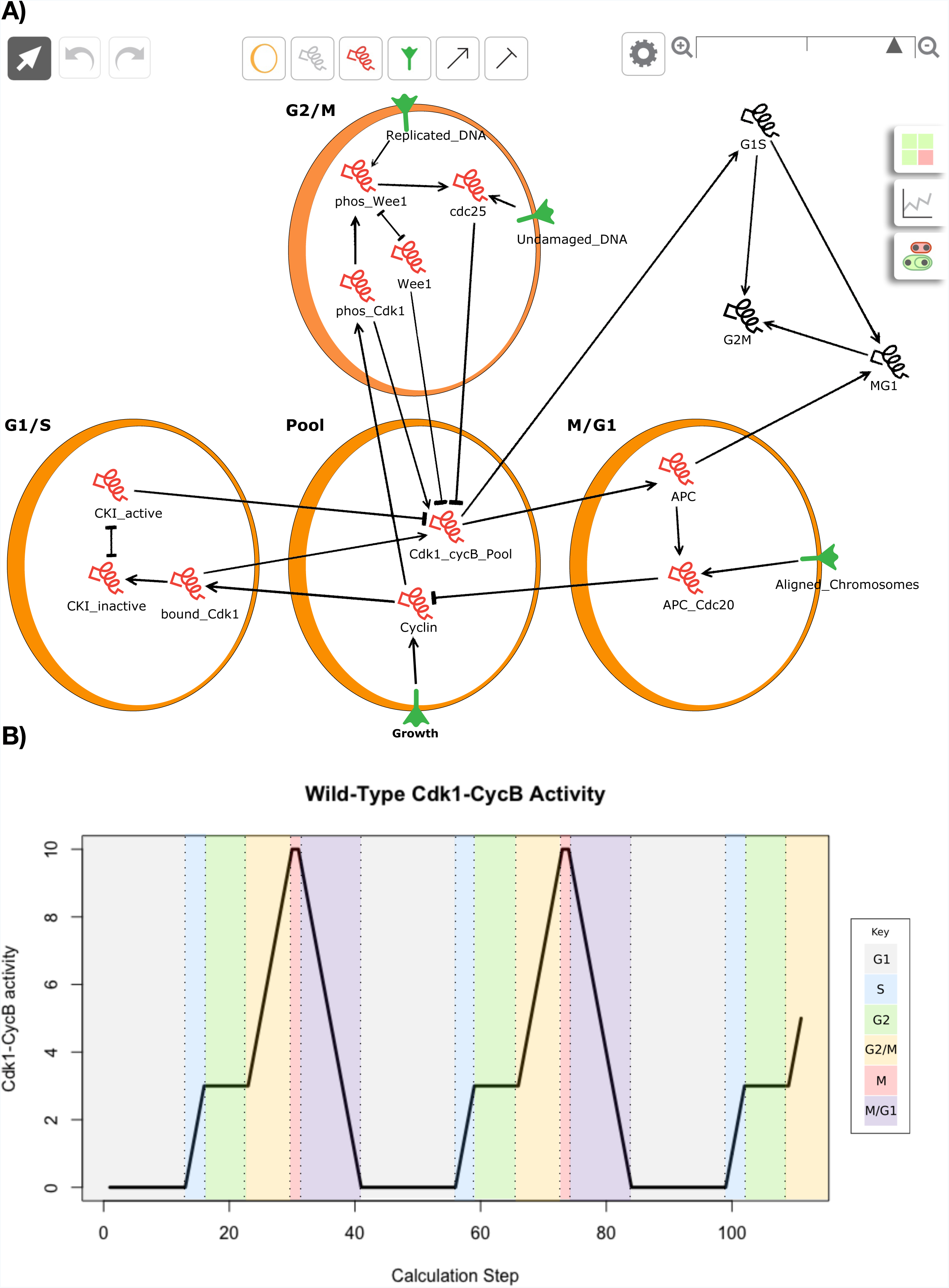
Qualitative Network of Eukaryotic Cell Cycle Regulation. **(A)** BMA Wiring diagram. The network is constructed around a central pool of the major cell cycle regulator cyclin dependent kinase (Cdk1) and its cyclin partner (cycB). This cell cycle transitions are triggered by changes in the Cdk1-cycB activity, which is regulated by a number of different components. CKI a cyclin kinase inhibitor and Wee1 kinase subunit inactive the Cdk1-cycB complex whereas the Cdc25 phosphatase activates the complex. Cdk1-cycB activity can also be destroyed via the Anaphase-promoting complex (APC) in combination with Cdc20, which target cyclin for degradation. The activities of the Cdk1-cycB activity can then be monitored by 3 extracellular markers; G1S, G2M and MG1. **(B)** BMA simulation of Cdk1-cycB activity. The solid black line indicates the progression of Cdk1-cycB levels through the cycle. Dotted lines and block colours represent distinct phases as determined by the key. The cycle repeats itself if growth conditions remain favourable, as is represented in this simulation.

## Pool Modulen

This module contains the “master molecules” of the cell cycle, that being cyclin-dependent kinases (Cdks) and their cyclin partner, which as the name suggests are required in order to activate the Cdks. Our model is limited to only a single Cdk-cyclin partnership, Cdk1-cycB for simplicity. This module is fuelled by the growth of cyclin levels which we assume can have unlimited binding capacity to Cdk1. Unlike cycB, intracellular Cdk1 concentration does not fluctuate throughout the cell cycle [27], we therefore model Cdk1 as being at a constant level which can accommodate the variations in cycB levels [20].

## G1/S Module

The G1/S module features mutual inhibition between Cdk1-cycB and CKI. This feedback loop is described as a “toggle switch” and is modelled using our BMA mutual inhibition target function. Here we model CKI as being present at high levels in G1 by assigning it an initial value of 10 (max based on our granularity choice). The input in this case is labelled as cyclin, which, as it increases causes an increase in bound_Cdk1 (i.e. heterodimer of CKI, Cdk1 & cycB) due to the initial high levels of CKI. As the CKI does not stop cyclin accumulation and binding to Cdk molecules, the rising Cdk1-cycB levels which are not opposed by CKIs soon tips the balance, phosphorylating the CKI and labelling them for degradation. The values chosen for the switch points can be found in **Supplementary table 2**.

## G2/M Module

Following the degradation of CKI and subsequent spike in Cdk1-cycB activity the cycle enters the G2/M module. This module features both mutual activation, between Cdk1-cycB and Cdc25, and mutual inhibition between Cdk1-cycB and Wee1 [19]. The later works in a similar way to that of Ck1- cycB and CKI, with a race occurring being between Cdk1-cycB and Wee1. The Cdk-cycB and Cdc25 mutual activation interaction on the other hand is a type of positive feedback loop, where Cdc25 and Cdk1-cycB activate each other rather than inhibit each other. This is modelled using our BMA mutual inhibition target function combined with the mutual activation target function. Here we model Wee1 as being present at high levels in G2/M by assigning it an initial value of 10. The input in this case comes from the G1_Cdk1 levels, which as it increases causes an increase in phos_Cdk1 (i.e. phosphorylated form of Cdk1-cycB) due to the initial high levels of Wee1. As Wee1 does not stop cyclin accumulation and binding to Cdk molecules, the rising Cdk1-cycB levels (which are not opposed by Wee1) soon tips the balance, phosphorylating the Wee1 and marking them as inactive. Inactive Wee1 maintains active Cdc25, thus decreasing Wee1 results in an increase in Cdc25 and thus the switch like activation of Cdk1-cycB. Again, the values chosen for the switch points can be found in **Supplementary table 2**.

## M/G1 Module

Once the Cdk1-cycB reaches a high level due to Cdc25 activation the cell enters mitosis. In order to exit this phase, the Cdk1-cycB activity must be destroyed and CKI levels stockpiled. This transition is aided by the Cdc20:APC complex, which itself is indirectly activated by Cdk1-cycB activity, causing degradation of CycB. This results in a substantial drop in Cdk1-cycB activity, which then allows CKI to rise again. This relationship is described as an oscillator based on a negative feedback loop, where Cdk1-cycB activates APC, which activates Cdc20, which then degrades CycB [19]. In the BMA, the negative feedback oscillators target function uses the default function. Therefore this was modelled simply by considering the whole cycle as a feedback loop by adding an inhibitory edge back to the cyclin B in order to create the desired response **(Supplementary table 2 & 3)**.

### Comparison to ODE Eukaryotic Cell Cycle Model Predictions

The initial conditions were set so that all nodes remained with an initial value of 0, except for CKI and Wee1 which are given an initial value of 10 (max based on our granularity choice). As Growth, Replicated_DNA, Undamaged_DNA and Aligned_Chromosomes are conditions that can be represented by a binary value, a value of 0 represents the absence of the cell phenotype, whereas a value of 1 corresponds to the presence of the phenotype. The initial values for all four of these phenotypes were therefore set to 1 to represent normal growth conditions. Simulation analysis, starting from this initial state leads to the initiation of a series of network states (ranging between 0 and 10 based on our granularity). These steps correspond to the biological time series of protein activation and inactivation that occur during the wild-type cell cycle (**Fig 3**, **B**).

Similar to the Tyson et al. [19] signal-response curve, **Fig 3**, **B** shows the cell progresses through the cycle via a number of steady states. Firstly, at low levels of Cdk1-cycB activity the cell will remain in G1. With increased growth it will eventually pass a critical point, resulting in the irreversible disappearance of G1. As the cell moves into the S phase the level of Cdk1-cycB continues to grow until it reaches an intermediate level (3 as determined by our target function). Here in the G2 phase the cell will continue to grow until it reaches the next critical threshold, where the G2 state will disappear. This gives rise to a large spike in Cdk1-cycB activity (driving the cell into mitosis) which then decreases as cycB is degraded by APC:Cdc20, signalling cell division and resetting the system for the next round of division. One added benefit of this model is its ability to continuously cycle, as highlighted in **Fig 3**, **B**.

### Simulation of Mutant Phenotypes Replicate Experimental Results Found in the Literature

In order to evaluate the accuracy of our model loss of function (KO) and over-expression (OP) mutations were carried out based on a sample of previous experiments found in the literature (**Table 1**). In our limited subset of mutant experiments 8 out of 9 cases were able to accurately replicate the experimental results found in the literature without making any modifications to the underlying model described above beyond modelling the mutations (**Fig 4**). For instance in the case of Cdc25 OP, studies in both yeast and mice have shown that over-production of Cdc25 result in premature entry into mitosis due to early activation of Cdk1-cycB [28,29]. In the in silico experiment, the same result can be discerned. Rather than needing 8 steps to pass through G2 the Cdc25 OP, the model only takes one step. Similarly less time is spent in M phase with only 1 step occurring versus 2 steps for wild types (WT). This results in the mutant model undergoing each cycle in fewer calculation steps, needing only 33 steps compared to the 41 needed in the WT model. Descriptions of the other seven successfully reproduced experiments can be found in **Table 1**. For the case concerning the CKI OP mutant, experimental results were not as clearly reproduced. Experimental evidence by Moreno & Nurse [30] showed that overexpression of Rum1, a fission yeast CKI, leads to delays in G1, with repeated S-phase and no M-phase. This is partially replicated in our mutant model, with there being a long delay in G1 phase (23 calculation steps compared to 14 in the WT model), as well as no M phase being reached (where Cdk1-cycB hits max value of 10). The model however still runs through the M/G1 phase rather than just repeating the S phase.

**Table 1:**
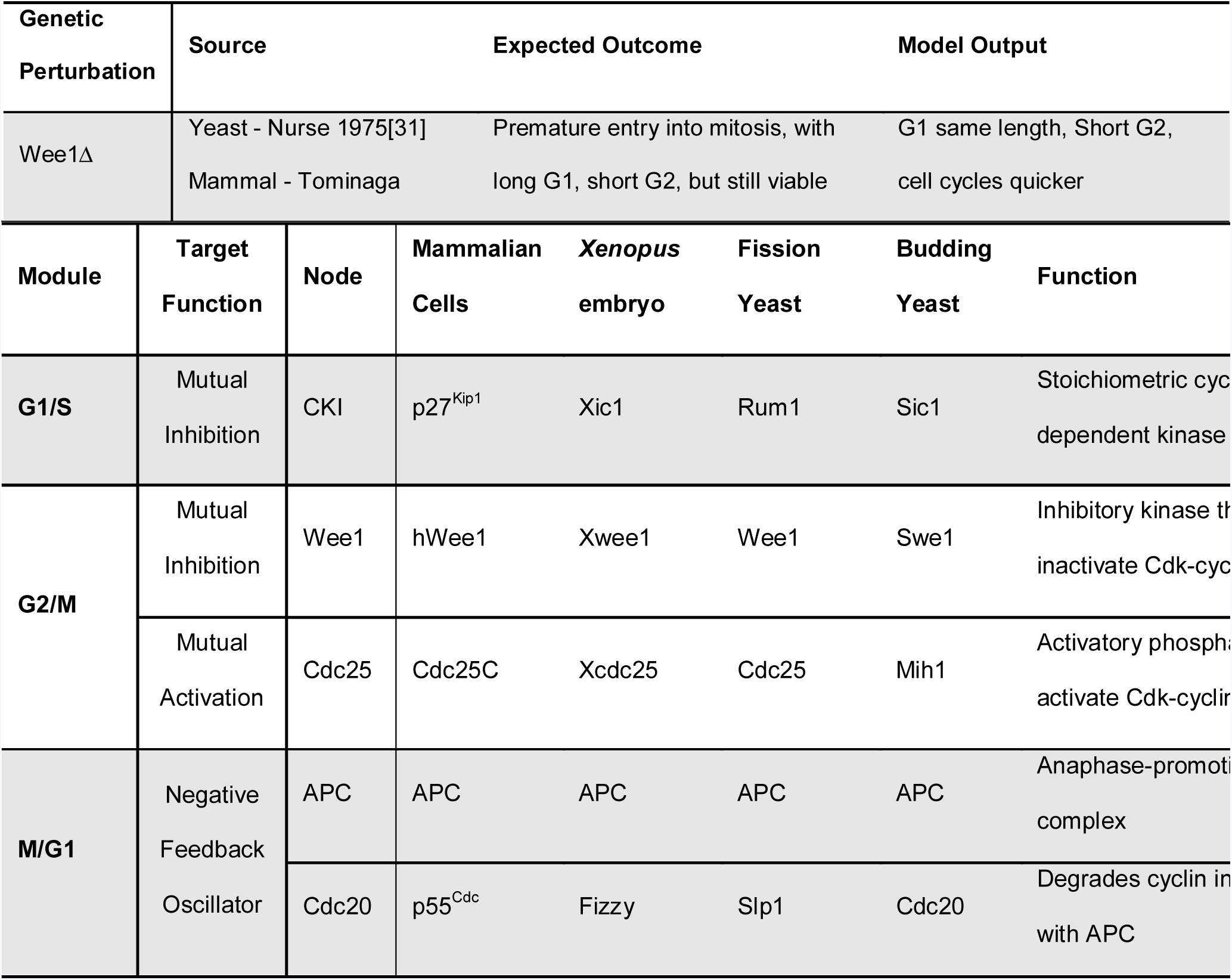
Mutant simulations reproduce described behaviour from the literature. Summary of experimental results are given in “Expected Outcome”, and in silico results are given in “Model Outcome”.

**Figure 4.**
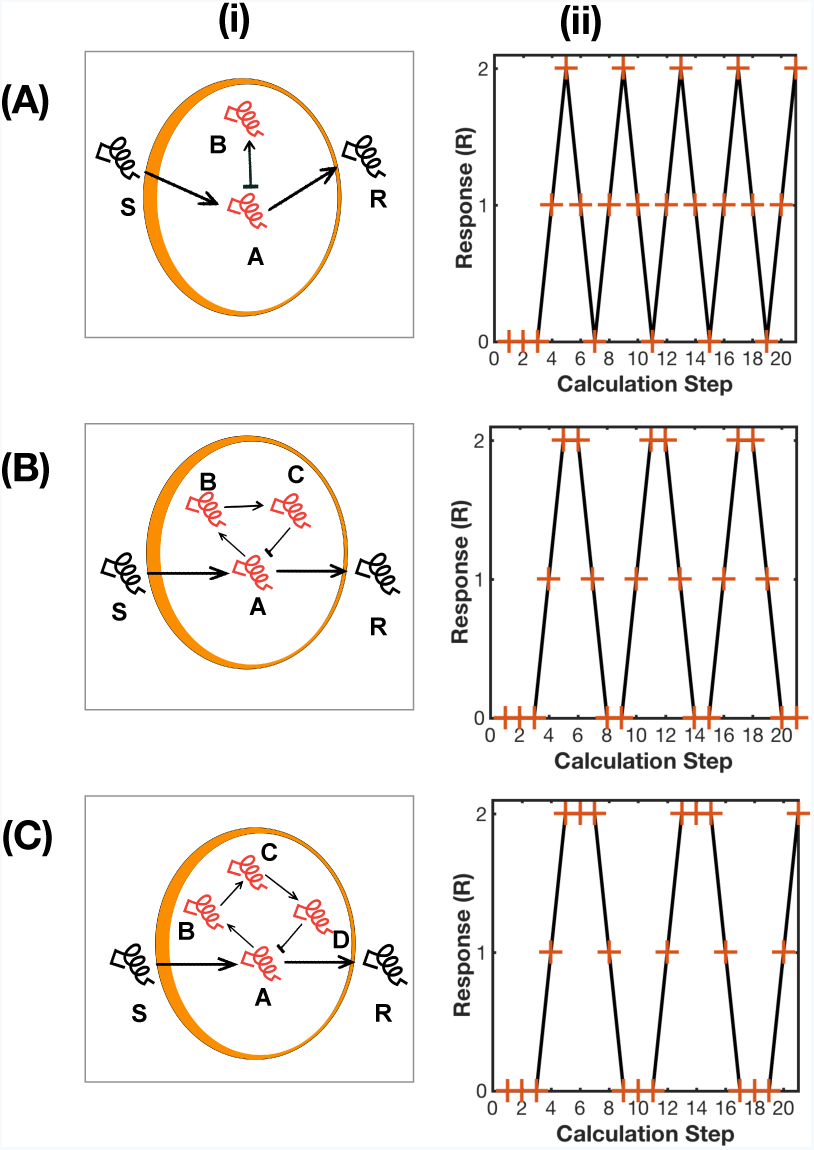
Mutant Phenotype Simulation Analysis. Depicts the temporal evolution of the network following perturbation of particular nodes. Each mutant perturbation can be compared to the wild type, which is listed first. Each distinct cell cycle phase is coloured coded according to the key provided. Each time step corresponds to each calculation step recoded in the BMA simulation which is exported as a CSV file.

## DISCUSSION

We present a library of novel Qualitative Network modules that can accurately replicate the biological behaviour of core, ubiquitous network motifs. We generate and compare our library based on biological behaviours defined previously [19], and confirm the modular nature of the library with the generation of a model for the eukaryotic cell cycle produced using motifs from the library. By simulating known genetic perturbations we further test this novel qualitative eukaryotic cell cycle model, highlighting its capacity to accurately replicate many well-known mutant phenotypes without the need for explicit parameterisation, as would generally be needed for ODE models. This study constitutes both a toolbox for biologists to construct elaborate networks with ease, but also an example of its application to a relevant biological system. The QN presented has much wider applications, with our working model having the potential to be adapted in order to provide much more dynamic details on the regulation of these core cell cycle components. Such a model could then be utilised to provide new insights into cell cycle regulation allowing the prediction of novel mutant phenotypes that have not been previously investigated. Not only could this provide a more thorough understanding of the underlying cell cycle regulatory principles, but also assist in the identification of a host of mutants that contribute to cancers or other pathologies, potentially allowing for the generation of novel drug therapeutics [40]. Similarly, in compiling this simple, easy to use BMA target function library we hope to encourage experimentalists to adopt this type of QN modelling as part of mainstream biological research. This would offer a wealth of advantages in terms of consolidating what is known about large networks into concise descriptions, as well as by allowing the generation of novel predictions about systems in the absence of large amounts of data and thus help focus experimental design.

Through the construction of our BMA motif/target function library we have been able to capture the dynamic behaviours of simple cell signalling pathways. Although networks can be modelled using ODEs, with behaviours being predicted using numerical simulations, this requires more complex and harder-to-obtain biological data and may equally appear mathematically complex to many biologists. As shown through the analysis of a model of eukaryotic cell cycle regulation, a relatively simple QN model can capture many of the advanced dynamic features of ODE models, including multistability and bifurcations. Simulation analysis of the described model shows strong similarities to that of the quantitative biological signal response curve, first proposed by Stern & Nurse [41], which was based on the results of multiple Cdk and cyclin knockout experimental studies. Like our model, they described the cycle as having three distinctive phases of Cdk activity, with the Cdk1-cycB levels transitioning through the cell cycle via different levels or bifurcations [22,41]. These levels or bifurcations are representative of firstly a stage of inactivity (G1) where Cdk activity remains low, secondly a stage of moderate Cdk activity sufficient to trigger S phase, and lastly a stage of high Cdk activity sufficient to initiate mitosis, all of which can be easily recognised in our model simulation [41]. This ability to model varying levels of Cdk activity sets our model apart from its Boolean counterparts, where only two levels of detail (“on” or “off”) can be captured. Through simple manipulation of our target functions we were also able to capture an extra layer of detail, by allowing our model to continue cycling over unceasing divisions when conditions remain favourable, a behaviour which has not always been replicated in previous studies [19,37,42]. The addition of this extra layer of complexity, showing sustained cell cycle oscillations, results in a model that is more representative of the true clock-like oscillatory nature of the cell cycle [43]. It is worth noting, however, that our model does not contain continuous, biologically measurable values for components, and as such is limited in its ability to interpret continuous experimental data.

As a means to further validate the model, loss of function and overexpression mutants were simulated, with the simplicity and generality of the model limiting the number of mutant phenotypes studies. Regardless of the simplification of using discrete modelling to represent continuous protein concentrations and interactions, the BMA model was capable of correctly modelling 8 out of 9 mutant phenotypes studied. All knockout mutations were correctly reproduced, with the model capturing dynamic properties such as phase length changes. For over-expression models, where the corresponding node range is set to max-max, 90% of the OP mutants studied corresponded accurately with experimental data, with CKI OP mutants being in partial agreement. This partial agreement is likely due to the minimalistic nature of our model, and could likely be overcome by using additional nodes to model the CKI interaction in more detail. Overall, the model produced using the BMA target function library accurately represents not only the WT regulation patterns of the general cell cycle control engine, but also the dynamic changes resulting from a number of mutants. This showcases our BMA target function library’s ability to be easily manipulated in order to model complex networks. Of particular note is the ability of the method to accurately generate protein behaviour through the simple addition of target functions from different modules that act on the same proteins, as is the case with the Cdk1-CycB node in our QN. This ability to draw together simple motifs to create realistic and useful biological networks demonstrates the validity of the approach and the opportunities that executable modelling makes available.

## METHODS

### Qualitative Networks

Qualitative networks (QNs) are an extension of Boolean models. In Boolean Networks, nodes are able to be in either and active (1), or inactive state (0), and are connected via functions that describe the mathematical relationship between them in an abstract way. Boolean Networks can be synchronous or asynchronous, that is − they may update every node simultaneously when a change is introduced in the system, or they can update in sequence from a propagation point. Qualitative networks are analogous to a synchronous Boolean Network, except that nodes are able to vary over a wide range of discrete values (called a granularity). Simple networks may be represented as Boolean, but Qualitative Networks may involve nodes with a greater range of values. For example, a node may have a range of 0-2 (granularity 3), where a value of 1 represents “normal activity” of an enzyme or gene product, and 0 and 2 represent low and high values respectively. This can be extended for much larger granularities, for example 0-10, where 10 represents maximal activity, and 0 represents minimal activity, with each discrete value in between representing a different concentration.

Nodes within a Qualitative Network are associated with either activatory or inhibitory relationships. Activatory relationships generally result in a response being high when a stimulus is high, and inhibitory relationships result in a response being low when a stimulus is high. Relationships between nodes are controlled by simple mathematical functions that describe the value that a node should represent, given its current inputs, and this function is called a target function. The values of nodes within a Qualitative Network are updated simultaneously when the network is simulated, and nodes will change their values in order to reach their target function gradually — changing value by only one each step. Due to the synchronous and defined nature of Qualitative Networks, they are deterministic, and susceptible to formal verification techniques. QN’s can stabilize and reach a single self-perpetuating state (called a stable point), but can also give rise to cycles and oscillations. Models and motifs described in this document are available in supplementary information and at https://github.com/shorthouse-mrc/biomodelanalyzer_targetfunctionlibrary.

### The BioModelAnalyzer (BMA) Platform

The BMA is an accessible, publicly available (https://www.biomodelanalyzer.org) graphical tool for discrete modelling and analysis of Qualitative Networks. The platform, with its user-friendly graphical interface, uses visual notations familiar to specialists in biology. BMA models are constructed on a gridded canvas upon which one or more cells, and cell elements (i.e. membrane receptor, cellular proteins etc.) can be placed and connected together with activatory or inhibitory links. To create a model, the user starts by dragging and dropping a cell onto the gridded canvas. These cells have no functional role in the analysis, being purely a visual aid to assist model design clarity. Cell elements are then placed in or outside of these cells, which can represent internal proteins, external proteins or membrane bound receptors. Connections between these cell elements can then be made using activatory arrows or inhibitory bar-arrows. Each cell element can then be labelled accordingly, using the simple drop down menus and a finite value range assigned, with the BMA default being [0,1], or Boolean. This range may be altered to add different levels of concentration, for example a range of [0,2] may represent “low”, “normal” and “high” concentrations of a protein or gene. If the user does not specify a target function for a node, then the BMA assigns a default target function. The default target function assigned within the BMA is described as:

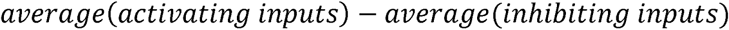

More complex target functions can be inserted for each node manually using an autocomplete function simplifying the use of correct syntax when referencing variables or using operators.

This underlying QN can then be analysed using simulation, stability analysis or Linear Temporal Logic tools each of which is accessible using the graphical interface. Simulation analysis shows the step-by-step execution of the model starting from a set point, based on either initial values specified by the user or a randomised start point. A graphical representation of all node values as they update over a user-defined number of time steps is produced, as well as a table of the simulation progression values, which can be exported as a CSV file for further analysis. Stability analysis can be used to test general properties of the model. If a model, given all possible starting conformations, will always result in a same self-perpetuating state, it is considered stable, and the graphical interface presents the user with the “stable values”. If stability is not achieved, however, the interface presents whether the system results in bifurcations (can potentially end in multiple states depending on the starting conformation) or oscillations (results in an infinite cycle). More advanced queries can be asked using the Linear Temporal Logic (LTL) interface, which allows the user to define simple or complex temporal logic queries with a drag and drop interface. LTL queries will return True, True sometimes, False, and False sometimes responses to queries, and the interface allows the user to see examples of systems where the behaviour occurs.

### Cell Cycle Model Generation

The model was composed of 3 main modules; G1/S, G2/M and M/G1 linked to a central node representing the level of Cdk1-cycB activity throughout the cycle. Each module was represented by a different cell in the BMA and labelled accordingly. The modules themselves were comprised of 6 key components namely; Cyclin, CKI, Wee1, Cdc25, APC and Cdc20, which regulate this Cdk1-cycB activity and thus the different cell cycle transitions (**Table 2**). These 6 components, modelled in their different chemical states (phosphorylated, active, inactive etc.) thus comprise a 20 node network, including 4 cell behaviours and 3 descriptive nodes linked by 28 interactions **(Supplementary Table 2 & 3)**. Three members of the BMA target function library were combined to create the cell cycle model, with the granularity set to 11 (A range of 0-10). This granularity, which differs from the default of 5 in our target function library, was chosen to accommodate the varying levels of Cdk1-cycB activity required, and to allow for clearer analysis of mutant phenotypes. Modules were initially generated based on the wiring and target functions from the BMA target function library examples, which were linked together through appropriate nodes **(Supplementary Figure 2)**. This method resulted in the creation of individual pools of Cdk1-cycB activity at the different cell cycle phases that fed into one central pool of Cdk1-cycB activity. To better represent the biological system, the model was then refined, by simply combining the Cdk1-cycB individual pool target functions into a single node via compound addition of each target function within the target function interface. To allow for multiple rounds of cell division, rather than the simulation of a single cell cycle, modification to the mutual activation target function was required. The mutual activation target function defines a one-way switch, and as such is not reversible. Here only nodes S and A (see figure 1, e, ii) and their associated target functions were used, thus allowing the cell to return from the high state achieved following the critical switch point activation.

**Table 2:**
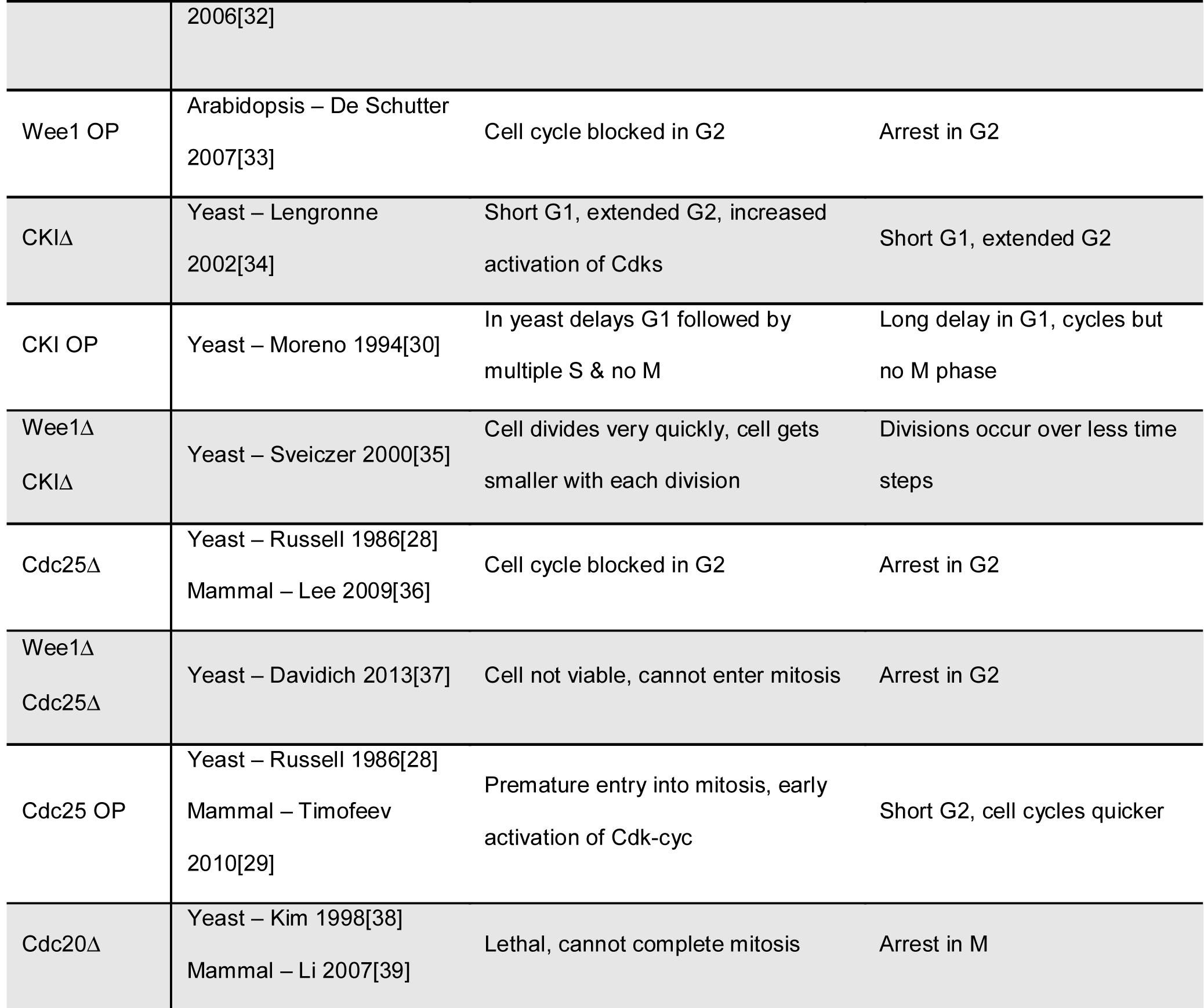
Cross-species nomenclature of key nodes within each module

### Knock-Out (KO) & Overexpression (OP) Analysis

In order to show if a model can faithfully reproduce known biological perturbations, loss of function and gain of function mutations can be analysed in BMA. A list of genetic perturbations curated from the literature is created and used to test the model. In the case of KO mutations, the corresponding node within the model range was set to a range of 0-0, corresponding to a permanently inactive state. OP mutations were simulated by setting the corresponding node range to max-max, (i.e. max based on the chosen granularity) simulating a permanently active state. Simulation analysis is then carried out, and the results compared to the wild-type simulation. Differences were then compared to known biological behaviours.

## ACKNOWLEDGEMENTS

We thank the Fisher and Hall groups for useful discussions. Work was supported by Medical Research Council core funding (D.S.) and Royal Society grant UF130039 (B.A.H).

## AUTHOR CONTRIBUTIONS

BH, CB, NP and JF conceived the study. DS and MP generated the target function library, YP developed the model of the cell cycle. DS and YP co-wrote the manuscript. All authors were responsible for editing of the manuscript.

## SUPPLEMENTARY INFORMATION

**Supplementary Figure 1.**
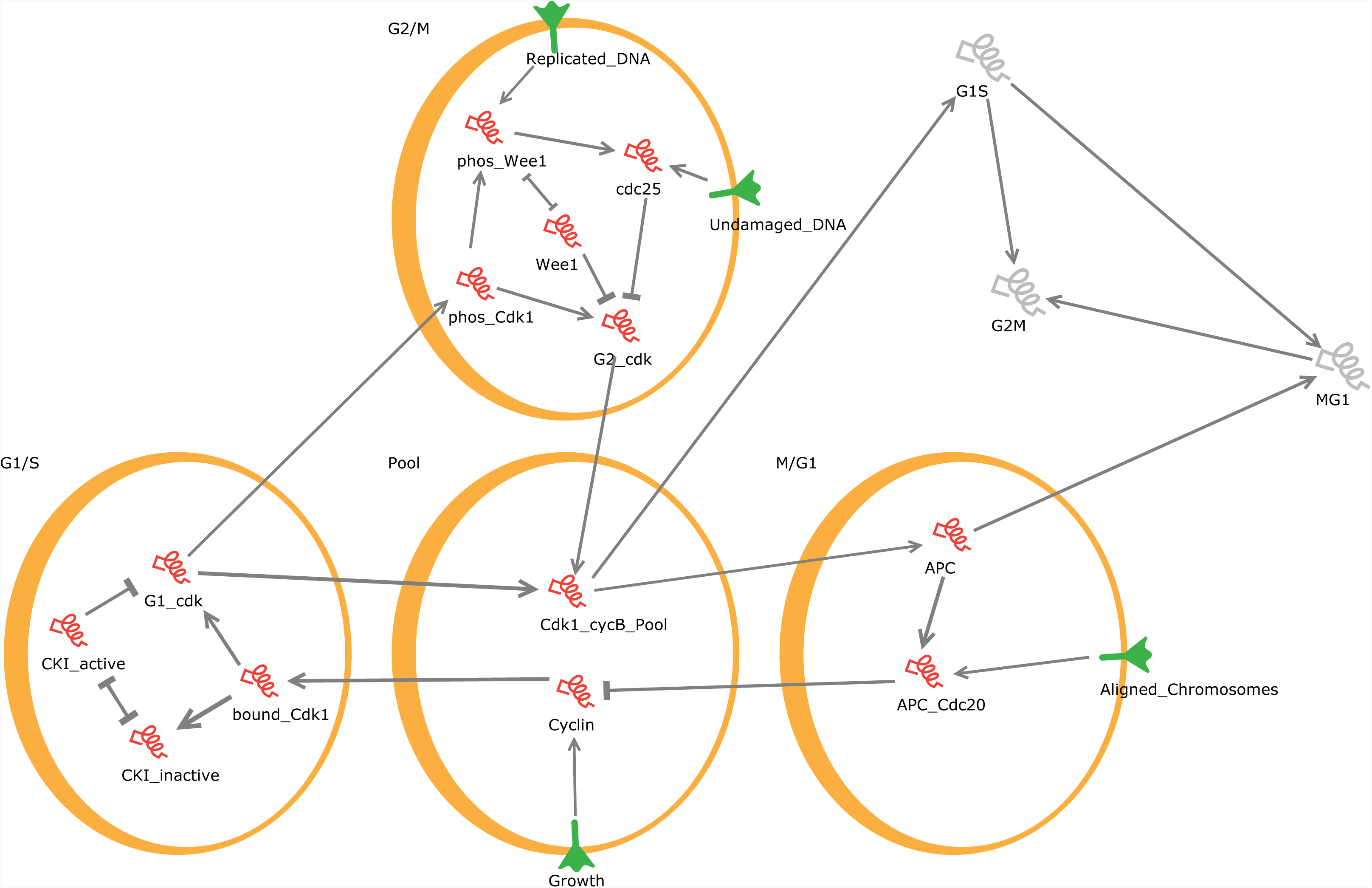
Adjustment of Negative Feedback Oscillatory Module. In this illustration, the rows correspond to **(A)** a one node system **(B)** a two node system and **(C)** three node system of negative feedback. The columns correspond to **(i)** the BMA wiring diagram translation and **(ii)** the BMA response curves. Each BMA wiring diagram contains a unique set of target functions located within particular nodes of the network which can be found in **Supplementary table 1**. S indicates the input Signal and R indicates the output Response with, in our case, letters A-D representing intermediate nodes. The graphs in (ii) are derived from simulation analysis carried out in the BMA. For all cases a simulation is run with a set signal input of 2 as an example, and the response output from the BMA simulation plotted based on the response per calculation time step. Comparison of the three different node length systems highlights that with increased number of nodes there is an increased length of oscillation response, as shown in the 20 calculation steps graphed, with the three node system **(C)** carrying out only 2 full oscillations compared to the one node system **(A)** which carries out 4 oscillations in the 20 calculation steps.

**Supplementary Figure 2.** Wiring Diagram. Wiring diagram composed using the network topology specified from the BMA target function library modules. Cdk-cycB activity is subdivided into individual pools which link to a central Cdk1-cycB pool. Subsequent models combine the target functions of the G1 Cdk-cycB activity and the G2-cdk-cycB activity together into one node which better represents the true biology.

**Supplemental Table 1.**
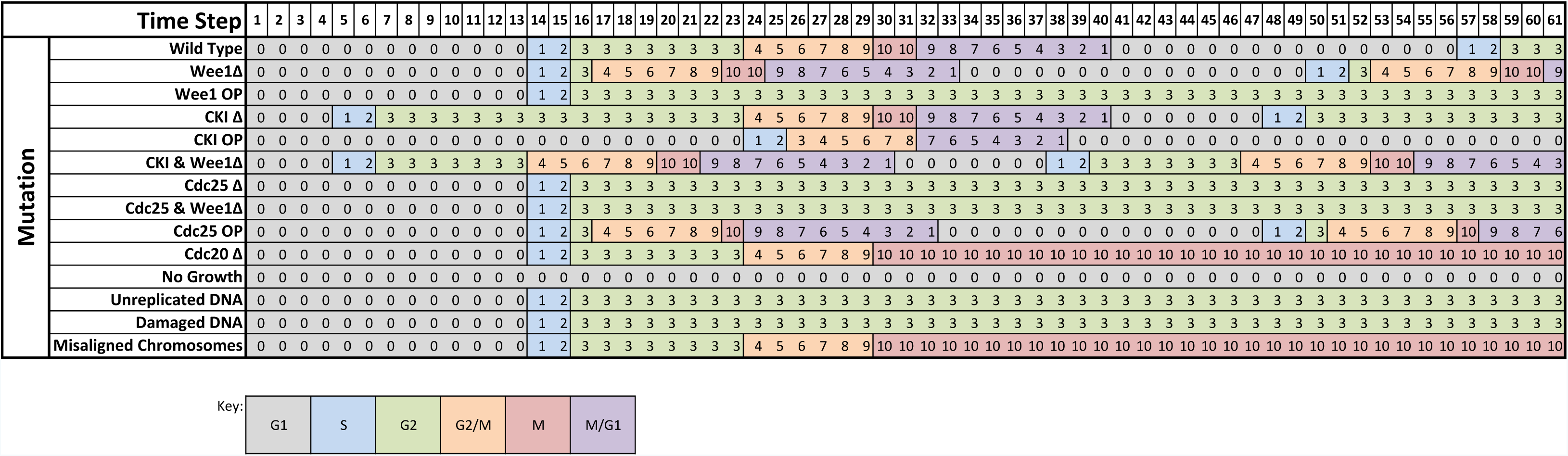
List of networks assessed in the manuscript. Paper reference refers to the figure in which the network occurs, included are the target functions for each node (if not the default), and comments. Included are the filenames and model names where the specific network can be found.

**Supplemental Table 2.** List of nodes in the network. Network ID refers to the internal label of the node. Full name is the common name found in the literature while Network Name is the name given in construction of model.

**Supplemental Table 3.** List of nodes in the network. Network ID refers to the internal label of the node. Full name is the common name found in the literature while Network Name is the name given in construction of model. The target function located in each node is found under the "Target Function" heading.

**Model Files**. We have additionally included all model files in .json format within an enclosed .zip file, they are also availabe at https://github.com/shorthouse-mrc/biomodelanalyzer_targetfunctionlibrary. Importing any file into the BMA will load the model and allow manipulation and simulation/stability analysis. Each file is named explicitly in Supplementary Table 1, with some files containing multiple models, which are referenced independently.

